# Self-consistent analytical solutions to the kinetics of lipid-induced protein aggregation

**DOI:** 10.1101/2025.07.10.664133

**Authors:** Alisdair Stevenson, David Voderholzer, Thomas C. T. Michaels

**Author notes:** These authors contributed equally (Alisdair Stevenson, David Voderholzer). Author to whom correspondence should be addressed. Electronic mail.

## Abstract

The aggregation of proteins into amyloid fibrils is a hallmark of several neurodegenerative disorders, including Parkinson’s disease. A growing body of experimental evidence highlights the significant role lipid membranes play in modulating this aggregation process, particularly for proteins such as *α*-synuclein. Despite this, there has been a lack of quantitative theoretical frameworks capable of describing the kinetics of lipid-induced protein aggregation. In this work, we develop an analytical model that explicitly incorporates lipid-mediated interactions into the aggregation kinetics. By formulating rate equations in terms of lipid surface coverage and applying a fixed-point analysis, we derive self-consistent solutions for the full timecourse of aggregation. Our model captures both one-step and two-step nucleation mechanisms and enables the prediction of key kinetic observables, including half-times and maximal growth rates. These results provide a quantitative foundation for interpreting experimental data and offer new mechanistic insights into how lipids influence the self-assembly of amyloidogenic proteins.

## I. INTRODUCTION

Polymerization of proteins and peptides into filamentous aggregates represents a fundamental form of self-assembly that is crucial for the functioning of biological systems, such as in the case of actin biofilaments^1^. However, aberrant filamentous protein aggregation forming amyloid fibrils is associated with severe clinical disorders such as Alzheimer’s and Parkinson’s disease^2–4^. Furthermore, due to their unique physicochemical properties, protein filaments are increasingly utilized as biomaterials in nanotechnology^5,6^. Given these diverse factors, the field of filamentous protein self-assembly has seen significant activity in recent years, and many efforts have been made to elucidate the underlying mechanisms of protein aggregation.

A fundamental question in the field of filamentous protein assembly has been to elucidate the microscopic mechanisms of the aggregation reaction. This information is crucial for the rational design of therapeutic strategies against protein aggregation disorders that target the aggregation process. Historically, mechanisms for simple chemical reactions have been studied through chemical kinetics by formulating rate laws and deriving analytical solutions, which link macroscopic reaction behavior to microscopic processes through quantitative comparison with experimental data. The application of these principles to the formation of protein filaments has yielded substantial insights, particularly through mathematical modeling approaches pioneered by Oosawa, who developed a framework for primary nucleation-driven aggregation^7^. This work was later extended by Eaton and Ferrone to include secondary pathways such as fragmentation and secondary nucleation^8,9^. More recent developments have provided analytical solutions to the complete aggregation kinetics with secondary pathways, representing a key step forward in linking theoretical models with experimental data, allowing quantitative comparison of theoretical predictions with experimental data describing protein aggregation kinetics^10–14^. However, one limitation of these chemical kinetics models of aggregation is that they typically assume spatial homogeneity, neglecting the impact of local environmental factors on aggregation behavior.

A particularly relevant example of a spatial heterogeneity that modulates aggregation is lipid surfaces, such as in the case of *α*-synuclein aggregation, a process associated with Parkinson’s disease. Here, the N-terminal region and part of the non-amyloid core of *α*-synuclein can bind to lipid surfaces and these interactions may form part of the *in vivo* function and pathology of *α*-synuclein^15–27^. Notably, lipid interactions have been shown to modulate the aggregation kinetics of *α*-synuclein, potentially altering nucleation and fibril growth pathways^23–44^. Here, we describe a kinetic framework that explicitly incorporates lipid-induced modulation of protein aggregation in terms of kinetic equations for the surface coverage of monomers and aggregates and derive analytical self-consistent solutions to these rate equations that describe the full timecourse of aggregation in terms of the underlying rate parameters. We discuss solutions in a number of limits, derive important characteristics of the reaction, including half-times, from first p rinciples. Our results provide a quantitative basis for interpreting experimental data and offer new insights into the impact of lipids on protein self-assembly, and were successfully applied to elucidate the mechanism of lipid-induced *α*-synuclein aggregation^45^.

The paper is structured as follows. In Section II, we introduce the theoretical framework of lipid-induced protein aggregation, developing kinetic equations based on surface coverage variables and considering both one-step and two-step primary nucleation pathways. Section III leverages the separation of timescales between lipid binding and aggregation to simplify the kinetic model and derive a reduced system within a slow manifold. In Section IV, we outline our fixed-point iterative approach for obtaining self-consistent analytical solutions, with Section V providing linearized solutions that serve as the initial approximation. Section VI presents first-order self-consistent solutions, while Section VII characterizes key features of the aggregation kinetics, including early-time behavior, steady-state yields, maximal growth rates, and half-time scaling laws. In Section VIII, we extend our model to account for lipid-limited conditions at low lipid-to-protein ratios.

## II. THEORY OF LIPID-INDUCED PROTEIN AGGREGATION KINETICS

We begin by introducing the kinetic equations for lipid-induced protein aggregation. We consider surface coverages of protein monomers and aggregates as the key quantities in our model, rather than their absolute concentrations (Fig. 1a). These surface coverages represent the fraction of the lipid surface covered by protein monomers or aggregates, taking values between 0 (empty surface) and 1 (fully covered surface), Fig. 1a. The protein monomer coverage *θ*_*m*_ is defined through the equation

**FIG. 1.**
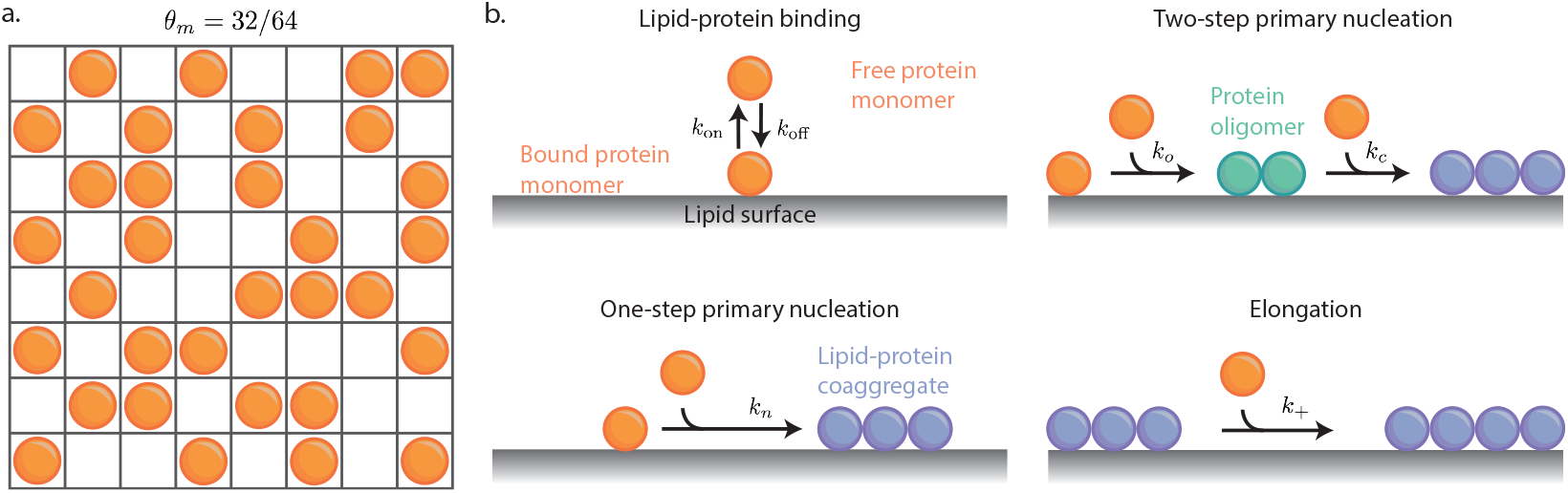
**(a)** Schematic representation of protein monomer surface coverage of a lipid surface. For simplicity, each protein monomer consumes one lipid binding site in this illustration (*β* = 1), but our theoretical framework can account for variable stoichiometries of protein-lipid binding through the *β* parameter. **(b)** Schematic representation of the elementary microscopic mechanisms underlying the formation and propagation of lipid-protein coaggregates considered in our model. These reaction mechanisms are both protein and lipid-dependent, where the variable contribution of bound and free protein monomers to coaggregate nucleation is described by the reaction orders *n*_1_ and *n*_2_. For elongation, bound, not free, protein monomers are required as the lipids are also incorporated into the coaggregate structure.

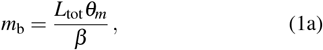

where *m*_b_ is the concentration of bound monomer, *L*_tot_ is the total lipid concentration (surface sites) and *β* is the average number of lipids that bind to a protein monomer in monomeric form. Similarly, we can introduce surface coverages for aggregates as

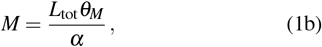

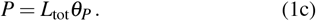

Here, we distinguish between fibril mass concentration *M* and fibril number concentration *P*. Consequently, *θ*_*P*_ is the probability of meeting a fibril end on the surface, produced by normalizing the number concentration of fibrils by the concentration of lipids. *α* is the average number of lipids that bind to a protein monomer in fibril form.

### A. Binding kinetics

We first consider the binding-dissociation kinetics of protein monomers to the lipid surface. To derive a dynamic equation in terms of protein monomer surface coverage, we describe the dynamics in terms of bulk concentrations of free and bound protein monomers to give

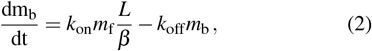

where *m*_b_ and *m*_f_ are the bound and free concentrations of protein monomer respectively, and *L* is the concentration of free lipid sites. The rate constants *k*_on_ and *k*_off_ describe the rates of protein-lipid binding and dissociation respectively (Fig. 1b). The bound protein concentration can be converted to surface coverage using Eq. (1a). The free protein monomer concentration can be described implicitly in terms of surface coverage terms by considering the conservation of protein mass in the system to give

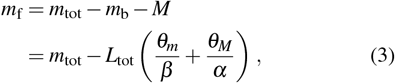

where the surface coverage of oligomers is neglected as *θ*_*S*_ *≪ θ*_*m*_ at early-times and *θ*_*S*_ *≪ θ*_*M*_ at late-times in the reaction. The fraction of free lipid sites is *θ*_*L*_ = 1 − *θ*_*m*_ − *θ*_*M*_ (again neglecting *θ*_*S*_), which follows from the conservation of lipid surface sites in the closed system, and can be converted to describe *L*:

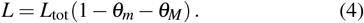

Taken together, we obtain the differential equation describing the dynamics of protein monomer surface coverage due to protein-lipid binding and dissociation as

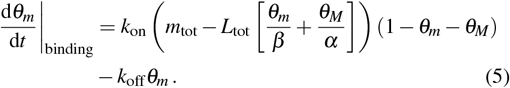

### B. One-step primary nucleation

In addition to the binding flux of protein monomers to the lipid surface, we consider fibril nucleation and growth events occurring in contact with the surface that produce lipid-protein coaggregates^37^, where these microscopic aggregation processes are shown graphically in Fig. 1b. In the simplest scenario, classical nucleation theory describes the nucleation process as occurring in one step, described by

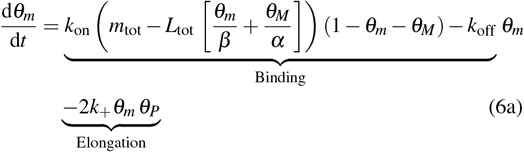

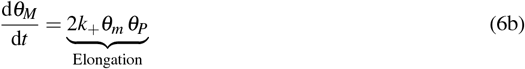

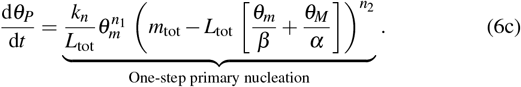

The first and second terms of Eq. (6a) describe the binding flux of protein monomers to the lipid surface, as described in Eq. (5). The third term of Eq. (6a) is the reaction flux for coaggregate elongation, where *k*_+_ is the rate of coaggregate elongation. This expression arises because of the assumption that coaggregate elongation is possible only via the incorporation of both protein monomers and lipids localized to the lipid surface. Therefore, the fibril elongation rate is proportional to *θ*_*m*_, rather than the concentration of protein monomers. The factor of 2 accounts for the possibility of adding monomers at either end of the fibrils. Eq. (6c) is the reaction flux of one-step primary nucleation, where *k*_*n*_ is the associated rate constant. We allow for the different possible contributions of free and bound protein monomers to the primary nucleation by using variable reaction orders *n*_1_ and *n*_2_ respectively. As fibril nucleation does not necessarily occur only on the lipid surface, immediately creating lipid-protein coaggregates, we convert the concentration to surface coverage with *θ*_*P*_ = *P*/*L*_tot_. Note that in Eq. (6a) and (6b), we have neglected the contribution of nucleation to the dynamics of *θ*_*m*_ and *θ*_*M*_. This is justified as the reaction flux due to filament elongation is significantly larger than the flux due to primary nucleation, which ensures the formation of sufficiently elongated structures. Consequently, elongation is the dominant reaction flux in the conversion process from proteins in monomeric to fibril form, rendering the dynamics of nucleation negligible for the dynamics of *θ*_*m*_ and *θ*_*M*_.

### C. Two-step primary nucleation

We also consider that the nucleation of filamentous structures often proceeds through a multi-stage process rather than a single step as proposed by classical nucleation theory. This non-classical nucleation pathway includes intermediate oligomeric states, where in our study, we examine a two-step primary nucleation mechanism where protein monomers first nucleate to form a generic, on-pathway, oligomer, that then converts to an elongation-capable coaggregate (Fig. 1b). A two-step primary nucleation mechanism has been identified in numerous amyloid-forming systems^46–48^, as well as in other nucleation processes, including crystallization^14,49–51^. To describe this scenario, we extend Eq. (6) to include the concentration of intermediate oligomers and additional microscopic mechanisms to give

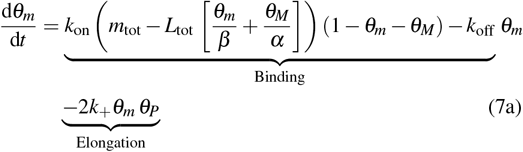

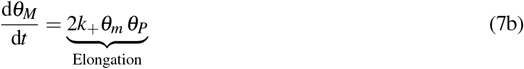

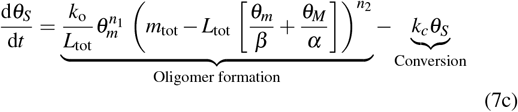

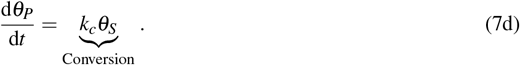

where *θ*_*S*_ is the concentration of oligomers and *k*_*o*_ and *k*_*c*_ are the rate constants for the primary nucleation of oligomers and conversion of oligomers to coaggregates respectively. Eq. (7a) and (7b) remain unchanged compared to the one-step model, Eq. (6). The first term in Eq. (7c) describes oligomer formation, with variable contributions of bound and free protein monomers. We implement a conversion process of oligomers to coaggregates with a linear dependence on the oligomer concentration, and no dependence on bound or free protein monomers, resulting in the oligomer conversion flux *k*_*c*_*θ*_*S*_, where *k*_*c*_ is the rate of conversion of oligomers to coaggregates. In principle, our model could describe a protein monomer dependent conversion process.

In Eq. (7), we assume that the process of oligomers binding to the lipid surface is not rate-limiting and thus do not explicitly describe it. Additionally, oligomer dissociation into constituent monomers is also disregarded as we do not have direct experimental measurements of oligomer concentration. In principle, our model can be extended to take both processes into account. If oligomer dissociation is explicitly described, the timescale for oligomer conversion is determined by an effective rate dependent on the rates of oligomer dissociation and conversion *k*_*e*_ = *k*_*d*_ + *k*_*c*_, thereby defining the steady-state timescale for oligomer concentration.

## III. PRE-EQUILIBRIUM OF PROTEIN TO LIPID BINDING

The first step to determine self-consistent solutions to the aggregation kinetics described by Eqs. (6) and (7) is to recognize the separation of timescales between the binding dynamics and aggregation reaction. Typically, the protein-lipid binding rate is much faster (on the scale of minutes) than the characteristic timescale of aggregation, which is the time for half the protein monomer mass to aggregate (on the scale of several hours), which has been validated experimentally^44^. With a separation of timescales between these processes, the kinetic models can be analyzed utilizing mathematical techniques based on matched asymptotics^52^ and allow for the division of the overall dynamics of the system into two stages:

- Initial layer – an initial rapid phase where aggregation is minimal, and a pre-equilibrium is established for protein-lipid binding.
- Slow manifold – a slower phase where aggregates form and grow, with the binding dynamics robustly maintaining a pre-equilibrium throughout.

Since the binding dynamics is much faster than the subsequent aggregation, we can neglect the initial layer and focus on the dynamics within the slow manifold when developing self-consistent analytical solutions to the aggregation kinetics (see Fig. 2 for a comparison of timescales). Here, the binding pre-equilibrium simplifies the dynamics to a lower-dimensional system, by setting the binding and unbinding terms in Eqs. (6a) and (7a) to zero due to the rapid preequilibrium dynamics, establishing a relationship between *θ*_*m*_ and *θ*_*M*_, described by

**FIG. 2.**
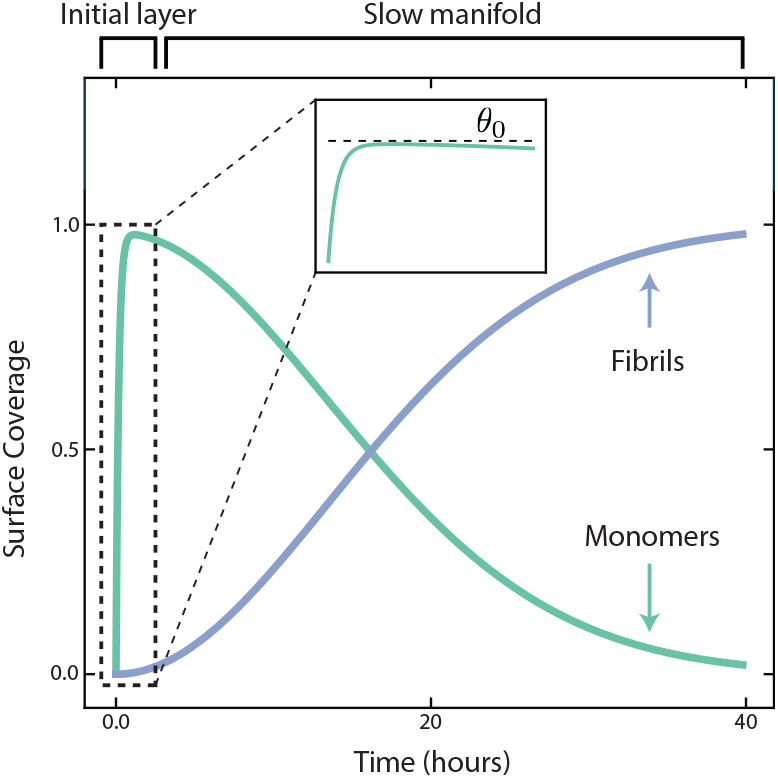
The separation of timescales between the kinetics of binding-dissociation of protein monomers to the lipid surface and subsequent lipid-induced protein aggregation reaction kinetics. Parameters used for this plot: *L*_tot_ = 100 *µM, m*_tot_ = 25 *µM, n*_1_ = *n*_2_ = 1, *β* = 28.2, *α* = 11.4, *K*_D_ = 3.8 *×* 10^−1^ *µM*^−1^, 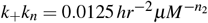.

**FIG. 3.**
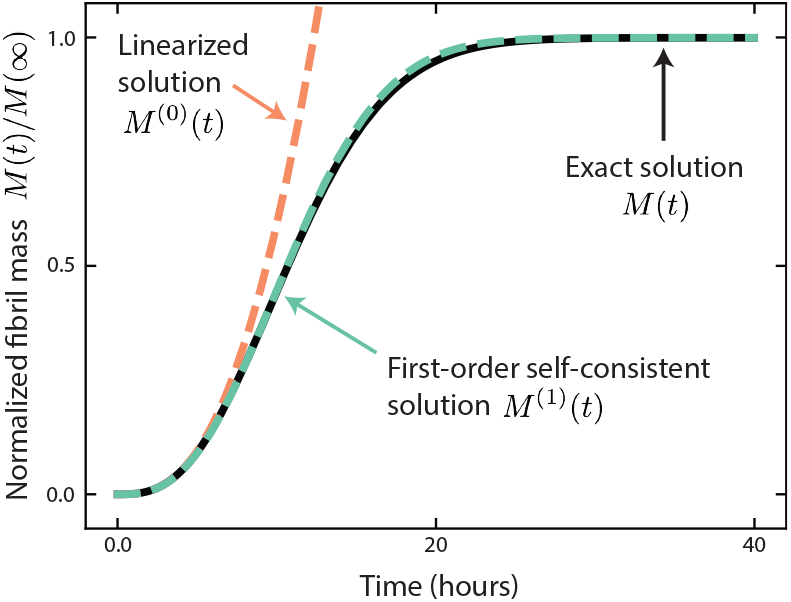
Comparison of the exact solution describing the dynamics of normalized fibril mass concentration for the two-step primary nucleation model derived from numerically evaluating the ODE system to the linearized analytical solution, *M*^(0)^(*t*), and self-consistent solution of order one, *M*^(1)^(*t*). Parameters used for this plot: *n*_1_ = 0, *n*_2_ = 1.5, *k*_+_*k*_o_ = 0.0118 *hr*^−2^ *µM*^−1^, *k*_c_ = 0.265 *hr*^−1^ and otherwise are identical to those used in Fig. 2.

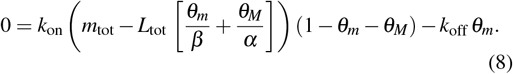

This equation can be solved for *θ*_*m*_ as a function of *θ*_*M*_ to yield

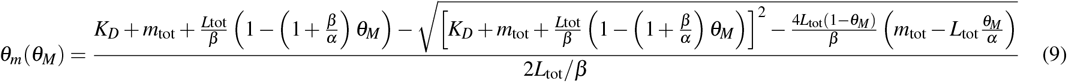

where 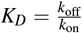 is the dissociation constant of protein-lipid binding. To make analytical progress, we linearize Eq. (9) on *θ*_*M*_ to yield the simplified expression

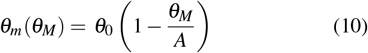

Where

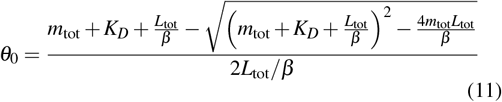

is the monomer surface coverage when *θ*_*M*_ = 0 and

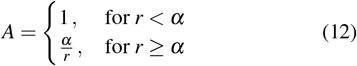

Where

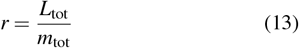

is the lipid-to-monomer ratio. The dynamics in the slow manifold are of a lower dimension due to the constraint, defined by Eq. (10), where *θ*_*m*_ can be described in terms of *θ*_*M*_ and reduce the number of independent kinetic equations. In particular, implementing the linearized binding pre-equilibirum, Eq. (10), into the kinetic equations of the one-step primary nucleation model, Eq. (6), produces a reduced set of dynamic equations in the slow manifold

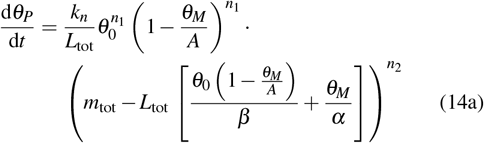

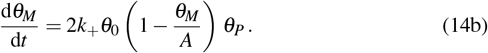

Similarly, implementing Eq. (10) into the two-step primary nucleation model, Eq. (7), produces

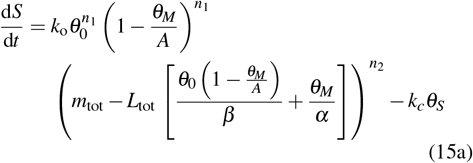

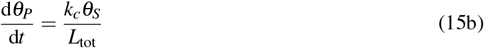

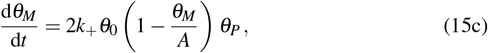

Therefore, by exploiting the separation of timescales between the binding dynamics and aggregation, we have reduced the system of kinetic equations to describe the dynamics in the slow manifold. In the following sections, we outline strategies based on fixed-point iterations to obtain accurate self-consistent solutions to these kinetic equations in the slow manifold.

## IV. SELF-CONSISTENT SOLUTIONS FOR LIPID-INDUCED AGGREGATION

The methodology for developing analytical solutions for Eqs. (14) and (15) is to utilize fixed-point iterations.^10,11^ This approach allows self-consistent solutions with increasing accuracy to be derived in an iterative process. Self-consistent techniques have been previously applied to derive analytical solutions for the kinetics of protein filament formation in the absence of lipids.^10,11^

To transform the differential equation systems (14) and (15) into a fixed-point problem, we integrate the kinetic equations such that they are of the form

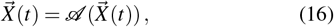

for some operator 𝒜 with 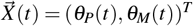 for the onestep primary nucleation and 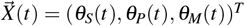 for the two-step primary nucleation model. Therefore, the desired solution to the system of differential equations (Eqs. (14) and (15)) is the fixed point 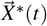 of the operator 𝒜. This problem is solved self-consistently if a fixed point 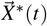 of the operator 𝒜 satisfying 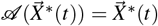 is found. According to the contraction mapping principle, for an initial guess 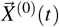 sufficiently close to the fixed point 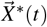, the fixed point can be computed iteratively as

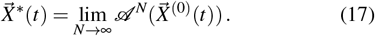

The convergence of the fixed-point iteration depends on the choice of the initial guess. Therefore, in the following section, we choose as an appropriate starting point for the iteration procedure the early-time linearized solution to the kinetic equations. This choice produces highly accurate self-consistent solutions for the full aggregation timecourse even after one iteration step.

To derive the fixed-point operators for the one-step and twostep models, we reformulate the kinetic equations Eqs. (14) and (15) as fixed-point problems through formal integration. For the one-step primary nucleation model, the fixed-point problem is:

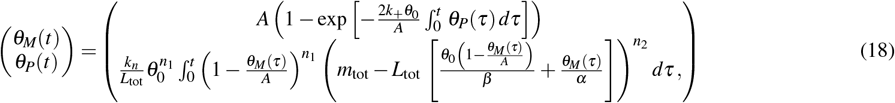

where the right-hand side of Eq. (18) is the fixed-point operator 𝒜. Similarly, Eq. (15) can now be rewritten as

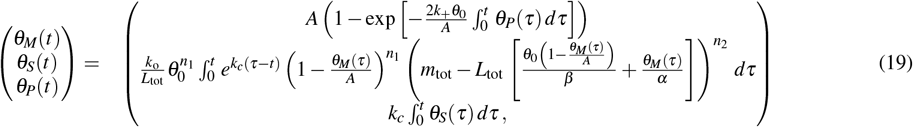

where the right-hand side of Eq. (19) is the respective fixedpoint operator 𝒜 for the two-step primary nucleation model.

Hence, the general form of *θ*_*M*_(*t*) in arbitrary order of the fixed point operation (except for zeroth order) will be

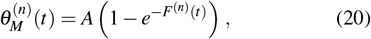

Where

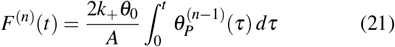

is a function that depends on the specific kinetic parameters in the model. Interestingly, the solution for 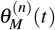 at any order in both the one-step and two-step models has this form. In the following sections we will derive explicit expressions for *F*^(*n*)^(*t*) through the application of the fixed-point iteration.

## V. SOLUTIONS TO THE LINEARIZED KINETIC EQUATIONS

As the starting point in our fixed-point analysis, we use the linear solutions to Eqs. (14) and (15) that emerge in the early-time limit, when the surface coverage of protein monomers can be assumed constant at its initial value. Mathematically, this is represented by setting *θ*_*m*_(*t*) equal to the initial value *θ*_*m*_(0) = *θ*_0_ for small times *t*, where *θ*_0_ is defined in Eq. (11). The solutions to the linearized kinetic equations capture the aggregation kinetics at early times, when monomer depletion through their incorporation into aggregates is negligible, and thus provide a valuable starting point for the fixed-point iteration method, facilitating its convergence.

Below, we discuss the solutions to the linearized kinetic equations for each model (one-step and two-step primary nucleation) separately.

### A. One-step primary nucleation model

Setting *θ*_*m*_(*t*) = *θ*_0_ in Eq. (14) and using *θ*_*M*_(*t*), *θ*_*P*_(*t*) *≪ θ*_0_ at early-times yields the following linearized kinetic equations

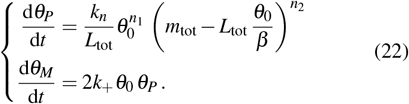

The solution for the initial values *θ*_*P*_(0) = *θ*_*M*_(0) = 0 reads:

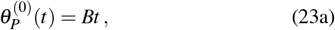

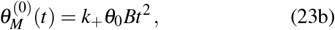

where 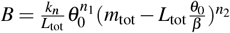.

### B. Two-step primary nucleation model

The linearized kinetic equations for the two-step primary nucleation model are:

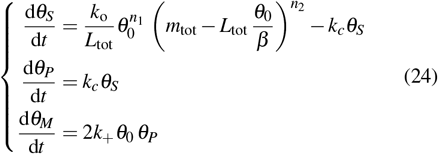

With the initial values *θ*_*S*_(0) = *θ*_*P*_(0) = *θ*_*M*_(0) = 0 the solution is

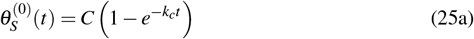

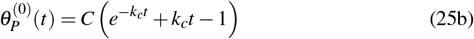

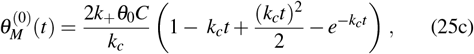

where 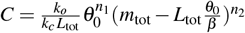.

## VI. SOLUTIONS TO THE NON-LINEAR MOMENT EQUATIONS

Accurate self-consistent solutions for the full timecourse of the aggregation reaction can now be obtained iteratively by applying the operator 𝒜 to the early-time solutions obtained in Sec. V.

### A. One-step primary nucleation model

The first-order self-consistent solution for the aggregate mass is obtained by substituting the linearized solution 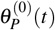, Eq. (23a), into the fixed point operator Eq. (18)

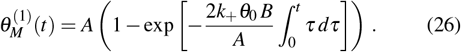

Performing the integration yields the first-order selfconsistent solution for *M*(*t*) as

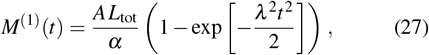

Where

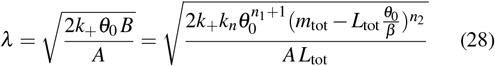

is an effective rate constant of aggregate proliferation through the combined effects of nucleation and elongation. This effective rate generalises the effective rate constant 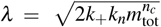 derived by Oosawa for nucleated polymerisation in the absence of lipid surfaces.

### B. Two-step primary nucleation model

Repeating the analysis for the two-step primary nucleation model, we obtain the first-order self-consistent solution 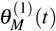 by inserting Eq. (25b) into the fixed-point operator Eq. (19)

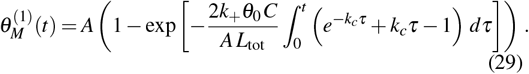

On integration, Eq. (29) becomes:

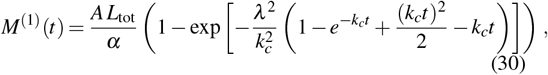

Where

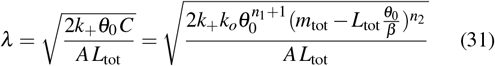

is the effective rate constant for aggregate proliferation, where *k*_*n*_ is replaced by *k*_*o*_. With these solutions, the effect of varying lipid and protein monomer concentrations on the aggregation kinetics is summarized in Fig. 4a and b, respectively.

**FIG. 4.**
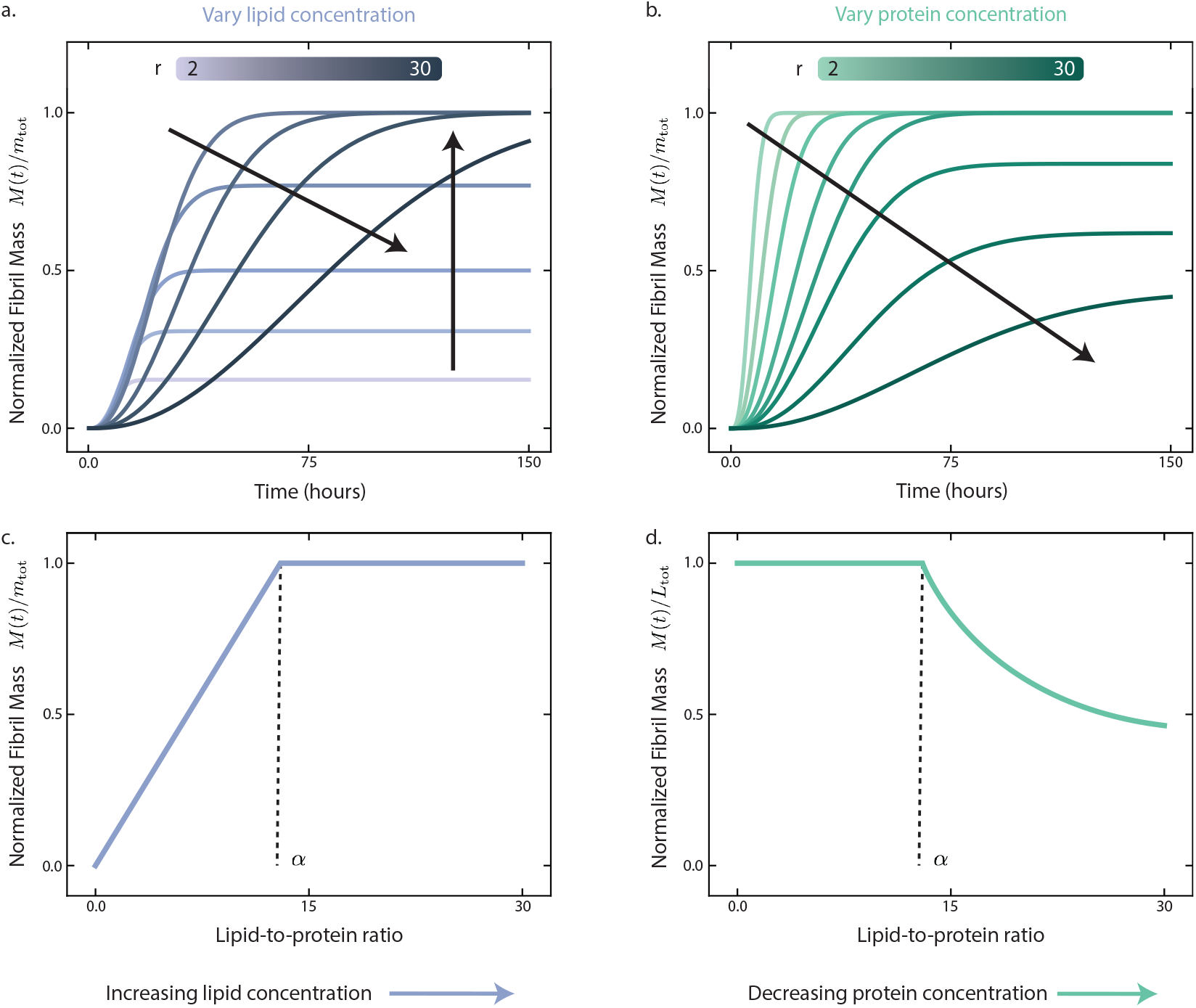
**(a) & (b)** Dynamics of normalized fibril mass predicted by the first order self-consistent solution to the two-step primary nucleation model for varying lipid-to-protein ratios, *r*, where the lipid **(a)** and protein **(b)** concentration is varied. **(a)** The vertical arrow indicates the change in behavior as lipid concentration is increased in the lipid-limited regime (*r* < *α*) and the diagonal arrow indicates the change in behavior as lipid concentration is increased in the protein-limited regime (*r* > *α*). **(b)** The diagonal arrow indicates the change in behavior as protein concentration is decreased across *r* values in both the lipid-limited and protein-limited regimes. **(c) & (d)** Dependence on the steadystate yield of lipid-protein coaggregates on the lipid-to-protein ratio, *r*, where the lipid **(c)** and protein **(d)** concentration is varied. Parameters used in this plot: *n*_1_ = 0, *n*_2_ = 1.5, *β* = 28.2, *α* = 13, *K*_D_ = 3.8 *×* 10^−1^ *µM*^−1^, *k*_+_*k*_o_ = 0.0118 *hr*^−2^ *µM*^−1^, *k*_c_ = 0.265 *hr*^−1^. Panel (a): *m*_tot_ = 20 and *L*_tot_ = 40, 80, 130, 200, 260, 400, 500, 600 *µM*. Panel (b): *L*_tot_ = 100*µM* and *m*_tot_ = 3.33, 4.76, 6.45, 7.69, 10, 15.38, 25, 50 *µM*.

## VII. CHARACTERISTICS OF LIPID-INDUCED PROTEIN AGGREGATION KINETICS

Having derived analytical expressions for the full timecourse of aggregation, we are now in a position to derive from first principles a series of important characteristics of lipid-induced protein aggregation. These include an analysis of the steady-state behavior as well as a discussion of half-time scaling.

### A. Steady-state behavior

The plateau concentration of aggregates can be found by taking the *t →* ∞ limit of Eqs. (27) and (30), yielding in both

Cases

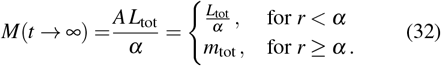

Interestingly, we find a biphasic thermodynamic behavior depending on the lipid-to-monomer ratio, where these solutions correspond to two possible outcomes of the aggregation reaction: *M*(*t* = ∞) = *L*_tot_/*α* or *M*(*t* = ∞) = *m*_tot_. In nonlinear dynamics theory, this behavior is defined as a transcritical bifurcation. When the lipid-to-monomer ratio *r* is below the critical value *α*, the final aggregate yield increases proportionally to the lipid concentration. Crucially, in this regime, not all protein monomers are converted into aggregates by the end of the reaction. There remains a surplus of protein monomers. By contrast, for *r ≥ α*, the final aggregate load is set by the available monomer concentration. In this case, all protein monomers are converted to fibrils, and there is an excess of lipids remaining at the end of the reaction. This biphasic behavior highlights the critical influence of the lipid-to-monomer ratio on the final state of the aggregation process. The steady-state behavior of the system when lipid and protein monomer concentrations are varied is summarized in Fig. 4c and d, respectively.

### B. Early-time limit

In the early-time limit *λt ≪* 1, the aggregate mass concentration follows a polynomial increase with time that corresponds to the solution to the linearized form of equation. For the one-step nucleation model, expanding the Eq. (27) for early times yields

**TABLE 1.** Comparison of the first-order self-consistent solutions to lipid-induced protein aggregation distinguishing the one-step and two-step nucleation models.

|  | One-step nucleation | Two-step nucleation |
| --- | --- | --- |
| Integrated rate law | $M^{(1)}(t) = \frac{AL_{\text{tot}}}{\alpha} \left( 1 - \exp \left[ -\frac{\lambda^2 t^2}{2} \right] \right)$ | $M^{(1)}(t) = \frac{AL_{\text{tot}}}{\alpha} \left( 1 - \exp \left[ -\frac{\lambda^2}{k_c^2} \left( 1 - e^{-k_c t} + \frac{(k_c t)^2}{2} - k_c t \right) \right] \right)$<br>$M^{(1)}(t) = \frac{AL_{\text{tot}}}{\alpha} \left( 1 - \exp \left[ -\frac{\mu^3 t^3}{3} \right] \right)$ (slow conversion) |
| Parameters | $\lambda = \left( \frac{2k_+ k_n \theta_0^{n_1+1} (m_{\text{tot}} - L_{\text{tot}} \frac{\theta_0}{\beta})^{n_2}}{AL_{\text{tot}}} \right)^{1/2}$<br>$A = \begin{cases} 1, & \text{for } r < \alpha \\ \frac{\alpha}{r}, & \text{for } r \geq \alpha, \end{cases}$ | $\mu = \left( \frac{2k_+ k_o k_c \theta_0^{n_1+1} (m_{\text{tot}} - L_{\text{tot}} \frac{\theta_0}{\beta})^{n_2}}{AL_{\text{tot}}} \right)^{1/3}$<br>$\theta_0 = \frac{m_{\text{tot}} + K_D + \frac{L_{\text{tot}}}{\beta} - \sqrt{(m_{\text{tot}} + K_D + \frac{L_{\text{tot}}}{\beta})^2 - \frac{4m_{\text{tot}} L_{\text{tot}}}{\beta}}}{2L_{\text{tot}}/\beta}, \quad K_D = \frac{k_{\text{off}}}{k_{\text{on}}}$ |
| Early-time limit | $M^{(1)}(t) \propto t^2$ | $M^{(1)}(t) \propto t^3$ |
| Half-time $t_{1/2}$ | $t_{1/2} = \frac{\sqrt{2 \log(2)}}{\lambda}$ | $t_{1/2} = \frac{\sqrt[3]{6 \log(2)}}{\mu}$ |
| Maximal growth rate $\sigma_{\text{max}}$ | $\sigma_{\text{max}} = \frac{AL_{\text{tot}}}{\alpha \sqrt{e}} \lambda$ | $\sigma_{\text{max}} = \frac{AL_{\text{tot}}}{\alpha} \left( \frac{2}{e^2} \right)^{1/3} \mu$ |

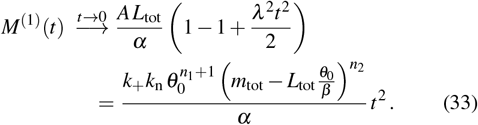

In this case, the early-time aggregate mass displays a quadratic dependence on time, *M*(*t*) ∝ *t*^2^. This result exhibits the same early-time behavior as the Oosawa model of nucleated polymerization, which also shows a quadratic increase at early times.

A similar polynomial early-time behavior is observed for the two-step nucleation model. Expanding 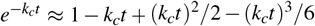 in Eq. (30) for *k*_*c*_*t ≪* 1 yields

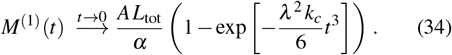

Expanding the exponential in Eq. (34) further for early times yields

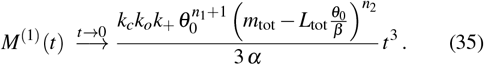

In the two-step nucleation model, the aggregate mass increases in proportion to *t*^3^, as opposed to the quadratic increase *t*^2^ observed in the one-step nucleation model. This reflects the additional complexity of the two-step nucleation process, where the formation of intermediates leads to a slower initial growth rate. More generally, the introduction of multi-step nucleation pathways can lead to even higher-order polynomial dependencies at early times. When nucleation involves multiple conversion steps before fibril elongation, the early-time kinetics are governed by:

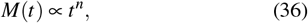

where *n* is the number of rate-limiting nucleation-conversion steps^53^. This means that higher-order nucleation cascades systematically increase the power-law exponent of the initial growth phase.

### C. Slow and fast conversion limits

In the limit of fast oligomer conversion, the argument of the exponential in Eq. (30) becomes

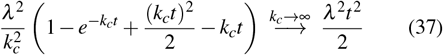

and therefore Eq. (30) recovers the first-order self-consistent solution for the one-step nucleation model

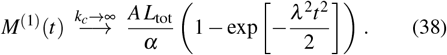

In the slow oligomer conversion limit, Eq. (30) simplifies to

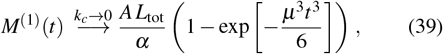

Where

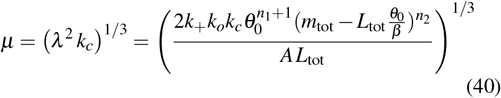

is a new effective rate controlling aggregate proliferation in the limit of slow conversion. Note that *µ* is the geometric average of the three rate constants of oligomer formation, oligomer conversion, and fibril growth.

### D. Maximal growth rate

The maximal growth rate quantifies the fastest rate of aggregate formation and is defined as:

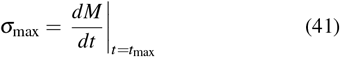

where *t*_max_ is the inflection point, defined by the equation 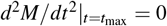 (see Fig.5a). For the one-step nucleation model, using Eq. (27), we find *t*_max_ = 1/*λ* and therefore the maximal growth rate is

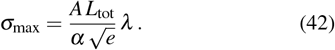

Similarly, for the two-step nucleation model, we obtain from Eq. (39) that *t*_max_ = 4^1/3^/*µ* and thus the maximal growth rate in this case reads

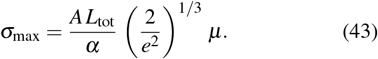

Fig. 5b presents a plot of the maximal growth rate *σ*_max_, revealing a peak at *r* = *α*, which marks the transition between the lipid-limited and protein-limited regimes.

**FIG. 5.**
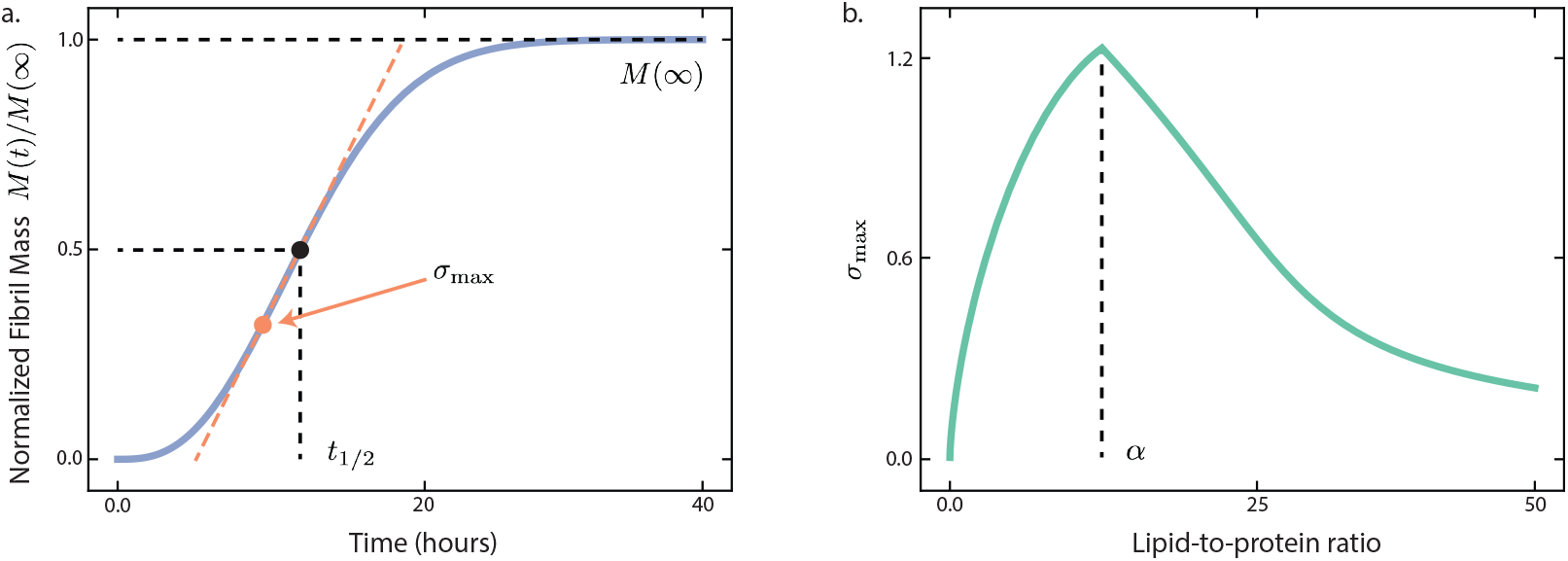
**(a)** Dynamics of normalized fibril mass predicted by the first order self-consistent solution to the two-step primary nucleation model detailing the location of maximal growth rate, *σ*_max_, and half-time of the aggregation reaction, *t*_1/2_. Parameters used for this plot: **(b)** Dependence on the amplitude of the maximal growth rate, *σ*_max_, on the lipid-to-protein ratio, *r*. The maximum of *σ*_max_ is achieved at the critical ratio *r* = *α* where the system switches from being lipid-limited to protein-limited with respect to coaggregate yield. Parameters used for this plot are identical to those used in Fig. 3.

### E. Half-time scaling

A key measure often used in characterising protein aggregation is the half-time *t*_1/2_, which is the time taken for the fibril mass concentration to reach half of its plateau value. In previous studies of homogeneous aggregation (i.e. in the absence of lipid surfaces), it has been shown that *t*_1/2_ displays a scaling relationship with the initial protein monomer concentration,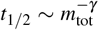.The scaling exponent, *γ*, depends on the reaction orders of the dominant nucleation mechanism. Therefore, experimental measurements of *γ* (e.g. through a double logarithmic plot of *t*_1/2_ with *m*_tot_) can reveal important mechanistic information about the underlying aggregation mechanisms. protein monomer concentration, *m*_tot_, and the initial lipid concentration, *L*_tot_ and examine the *t*_1/2_ scaling. Interestingly, we find a distinct *t*_1/2_ scaling when varying *m*_tot_ or *L*_tot_ and, for both of these cases, the scaling of *t*_1/2_ exhibits multiphasic behavior with distinct scaling exponents in the lipid and protein limited regimes. For this reason, we introduce two scaling exponents *γ*_*L*_ and *γ*_*p*_ to capture the dependence of *t*_1/2_ on *L*_tot_ and *m*_tot_, respectively:

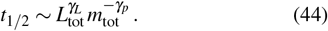

We now derive explicit expressions for these scaling exponents *γ*_*L*_ and *γ*_*p*_ valid in the different regimes.

#### 1. Fast conversion limit (one-step nucleation)

The half-time of aggregation can be found explicitly by the solving 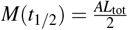, which using Eq. (27) yields

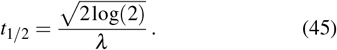

It is convenient to explicitly consider the form of Eq. (45) in the two regimes *r* < *α* and *r ≥ α*:

- When *r* < *α*, we have *A* = 1 and Eq. (45) reads explicitly

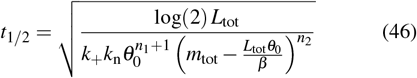
- When *r ≥ α*, we have *A* = *α*/*r* and therefore Eq. (45) becomes:

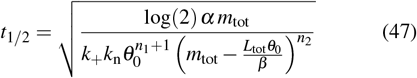

To understand how Eqs. (46) and (47) scale in the limit of low/high values of *m*_tot_ and *L*_tot_, four cases were examined.
- **Case 1:** *m*_tot_ = constant & *L*_tot_ *→* 0 (*r* < *α*). In this limit, *θ*_0_ *→ m*_tot_/(*m*_tot_ + *K*_*D*_). Thus, using Eq. (46) in the lipid limited regime, we find

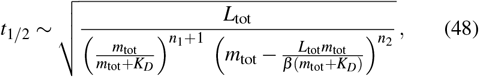

and as 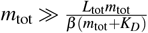 and *m*_tot_ = constant, we arrive at:

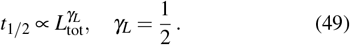
- **Case 2:** *m*_tot_ = constant & *L*_tot_ *→* ∞ (*r* > *α*). In this limit, 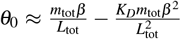 The *t*_½_ scaling can now be found by examining (47):

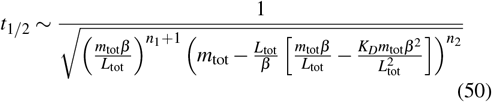

so as *L*_tot_ *→* ∞:

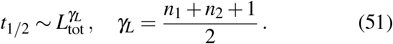
- **Case 3:** *L*_tot_ = constant & *m*_tot_ *→* 0 (*r* > *α*). In this limit, 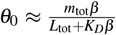 and using Eq. (47) in the protein limited regime, we find:

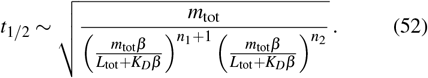

In other words

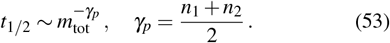
- **Case 4:** *L*_tot_ = constant & *m*_tot_ *→* ∞ (*r* < *α*). In this limit, *θ*_0_ *≈* 1 and therefore from Eq. (46), we find

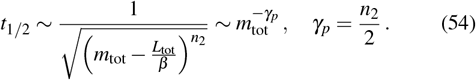

The scaling exponents can be converted to the form *t*_1/2_, and the scaling results are summarized in Fig. 6, where the scaling behavior of the one-step primary nucleation model is identical to the scaling of the two-step primary nucleation model with fast oligomer conversion.

**FIG. 6.**
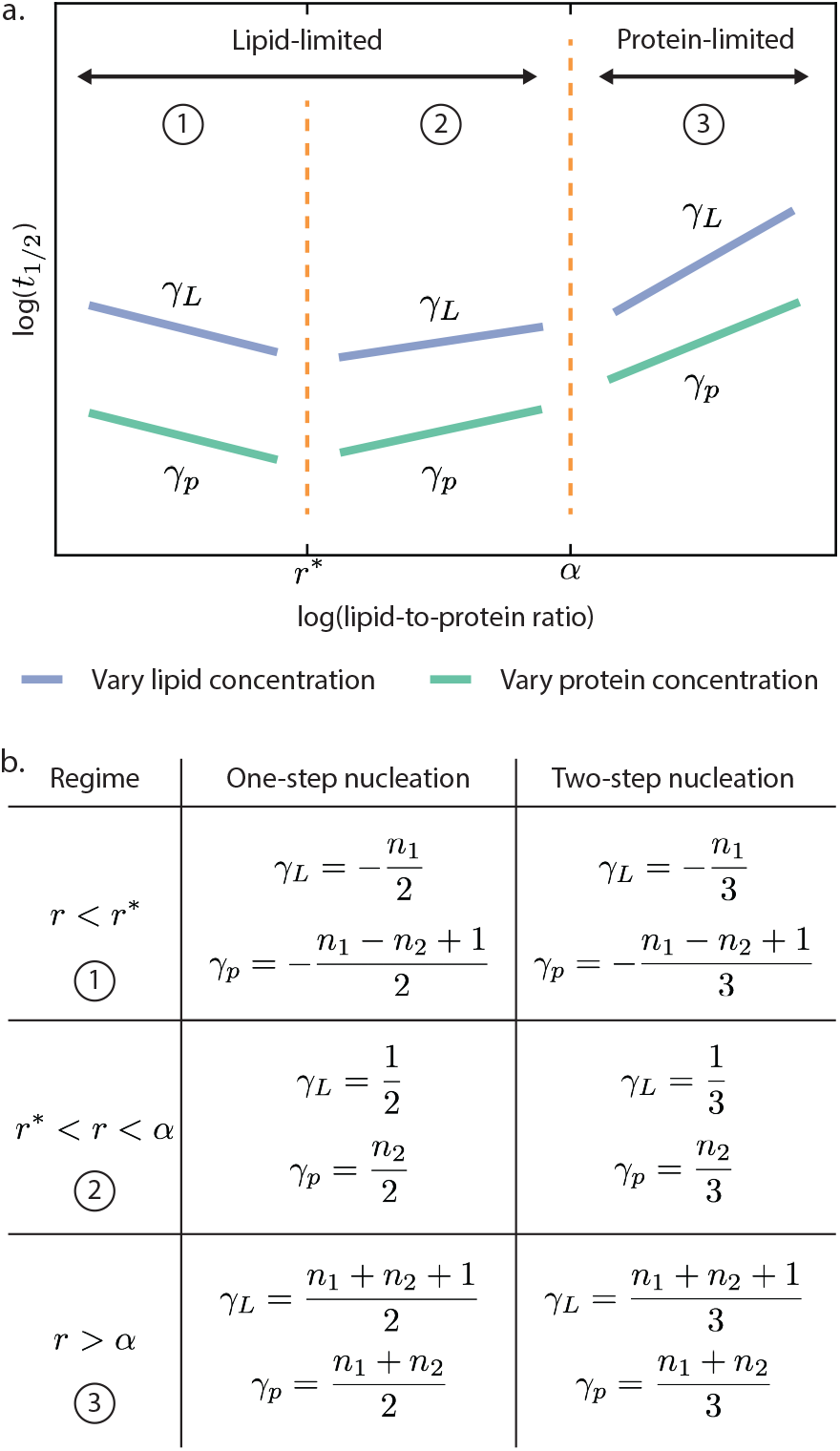
**(a)** Schematic log-log plot of *t*_1/2_ against *r* qualitatively describing the six distinct scaling regimes, contingent upon whether the aggregation reaction is limited by initial lipid or protein concentration. In region 1, the reaction is lipid-limited and *r* < *r*^*^. In region 2, the reaction is lipid-limited but *r*^*^ < *r* < *α*. In region 3, the reaction is protein-limited as *r* > *α*. **(b)** Table summarizing theoretical predictions for the scaling exponents *γ*_L_ and *γ*_p_ depending on *r*.

#### 2. Slow conversion limit (two-step nucleation)

In the limit of slow oligomer conversion, we use Eq. (39) to determine the half-time, yielding the following expression:

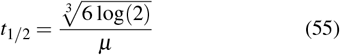

Again, it is useful to distinguish between the two regimes *r* < *α*

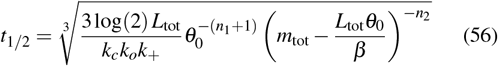

and *r* ≥ *α*, respectively

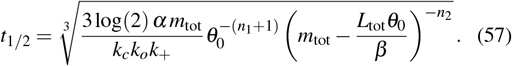

By using the same limits for *θ*_0_ as in Section VII E 1, we find the following scaling exponents for *t*_1/2_. In the regime *r* < *α*,

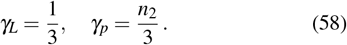

In the regime *r* > *α*, we find

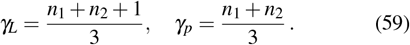

## VII. LOW LIPID-TO-MONOMER RATIO: LIPID-TRANSPORT LIMITED REGIME

Experimental data indicate that when *r* falls below a critical threshold (*r*^*\**^), kinetic traces slow down as *r* decreases. This behavior suggests that the elongation process, which is typically lipid-driven, becomes constrained by lipid scarcity, requiring a revision to our theoretical framework. Rather than proceeding as a single-step reaction, lipid-driven elongation transitions to a two-step process at low *r*. To account for this, we introduce a correction factor that modifies the nucleation and elongation fluxes in our equations. The key adjustment involves correcting *θ*_*m*_ in Eq. (6), respectively, Eq. (7) with a term that explicitly accounts for lipid scarcity

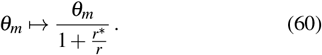

This factor ensures that, for *r* < *r*^*\**^, the effective lipiddependent aggregation rate scales with *r*/*r*^*\**^, reflecting the limited availability of lipid binding sites. In contrast, for *r* > *r*^*\**^, the correction term approaches unity, preserving the original form of the model.

### A. Self-consistent solutions

With revised nucleation and elongation fluxes, self-consistent solutions retain their original structure (Eq. (30)), but incorporate effective scaling terms for *λ* and *µ*. Specifically, we introduce the following transformations:

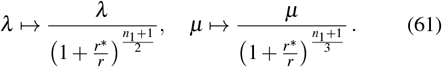

These modifications ensure that the lipid limitation is explicitly accounted for in the rate equations, leading to corrected aggregation kinetics in the low-*r* regime.

### B. Half-time scaling

The introduction of the lipid-dependent correction in *λ* and *µ* directly impacts the scaling behavior of the half-time (Fig. 6). Specifically, for *r* < *r*^*\**^, the scaling exponents *γ*_*L*_ and *γ*_*p*_ are reduced due to the lipid limitation. For one-step nucleation, the correction modifies the scaling exponent as follows:

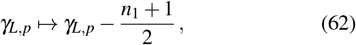

while for two-step nucleation, the reduction follows a different scaling factor:

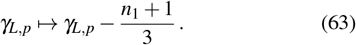

## IX. CONCLUSIONS

In this work, we have developed a self-consistent theoretical framework to describe the kinetics of lipid-induced protein aggregation. By explicitly incorporating lipid-dependent nucleation and elongation processes, our model extends classical polymerization theories^7,10^ to account for the critical role of lipid availability in modulating aggregation behavior. A key advantage of this framework is the availability of analytical solutions, which will enable the interpretation of experimental aggregation data in terms of the underlying rate constants of the microscopic steps of aggregation. By offering a quantitative means to extract kinetic parameters from experimental data, this framework lays the groundwork for future studies exploring lipid-mediated protein aggregation in physiological and pathological settings. Future extensions of this model could incorporate additional processes such as secondary nucleation and aggregation inhibitors to further refine our understanding of lipid-protein interactions in aggregation.

## ACKNOWLEDGMENTS

This work was funded by ETH Zurich (AS, TCTM) and the Swiss National Science Foundation (grant no SNS 219703 to TCTM).

## DATA AVAILABILITY

The data that support the findings of this study are available within the article.

